# Chemical and technological characteristics of sugarcane as a function of pelletized organomineral fertilizer with filter cake or sewage sludge sources

**DOI:** 10.1101/2020.07.16.206136

**Authors:** Carlos André Gonçalves, Reginaldo de Camargo, Robson Thiago Xavier de Sousa, Narcisa Silva Soares, Roberta Camargos de Oliveira, Mayara Cristina Stanger, Regina Maria Quintão Lana, Ernane Miranda Lemes

## Abstract

Sugarcane is one of the major alternatives for the biofuel sector and its large production has considerable environmental impacts. Organomineral fertilizers formulated with environmental liabilities such as filter cake and sewage sludge positively affect parameters of plant growth and development. The objective of this study was to evaluate the chemical and technological characteristics of sugarcane fertilized with pelletized organomineral fertilizers based on filter cake and sewage sludge. Eight field treatments were studied, being three levels of organomineral (50, 100, 150%) of two organic matter sources (filter cake, sewage sludge), plus a control with 100% of the recommended fertilization via mineral fertilizer and no fertilization control (0%). Sugarcane was evaluated during two consecutive harvest, and the amount of stalks per hectare (ton ha^−1^), the sugarcane productivity (ton ha^−1^), the quantity of sugar per hectare (TSH, ton ha^−1^), and the chemical and technological analysis of the sugarcane juice: pol (%), brix (%), purity (%) and fiber (%) were evaluated. None of the organomineral sources or doses differed from the exclusive mineral fertilization. The analysis of both harvests indicated that the first cut was the most productive since the responses of the first harvest were superior or similar to the second harvest. The recommended organomineral dose to obtaining maximum quantitative and qualitative sugarcane results was between 102 and 109% of the regular recommendation for mineral fertilization, regardless of the organic source in the first sugarcane harvest. In the second sugarcane harvest, sewage sludge source increase by 4.68 and 4.19% the total amount of sugar per hectare and the quantity of sugarcane compared to the sugarcane filter cake source. Sewage sludge and sugarcane filter cake as sources for organominerals are viable alternatives and advantageous in economic and environmental terms for the cultivation of sugarcane.

## Introduction

The replacement of fossil fuels by renewable, economically viable and less polluting sources became a need and a feasible option in the past decades. This was especially due to the use of ethanol fuel on a large scale obtained mainly from sugarcane and maize. Brazil and the United States are countries that began to invest in this type of biofuel heavily. Focusing on ethanol from maize, the United States became its largest producer, with more than 60 billion liters per year [1]; Brazil, in 2018/19, reached the record production of 33.14 billion liters of ethanol mainly from sugarcane, being maize responsible for the generation of only 1.35 billion liters [2].

Sugarcane is considered one of the major alternatives for the biofuel sector, not just due to the high potential for ethanol production, but also for their respective byproducts. The production mills have sought to improve the efficiency of ethanol production, increasing its supply and reducing production costs. Concomitantly, the ethanol industry seeks the use of sustainable agricultural inputs and eco-friendly cultivation techniques. The organomineral fertilizers are a good example of these improved techniques. The great efficiency demonstrated by these fertilizers in many crops gradually conquer the market. In addition to the possibility of aggregate new technologies to the product, the organominerals also make use of environmental liabilities. These materials can contaminate the environment when not receiving an environmentally correct destination.

The organomineral fertilizers in Brazil must meet the Normative Instruction n°. 25, from 23 July 2009 of the Brazilian Ministry of Agriculture, Livestock and Food Supply [3], requiring the assisted composting of the organic matter used; the minimum guarantees of organic carbon (8%) corresponded to a content of approximately 40% of composted organic matter, and 60% of mineral fertilizer, plus the nutrients declared on the label and based on soluble and total levels.

The sugarcane filter cake is retained in rotary filters after sugarcane processing in industries and has been the main organic residue in the composition of organomineral fertilizers. Filter cakes were initially obtained only in the process of sugar production, but currently, the alcohol mills are producing considerable amounts of filter cake [4]. Good results of fertilization are being observed when filter cakes are applied to the soil (directly, or as organomineral), generating positive responses to the productivity of crops such as sugarcane [4-6]. Additionally, sugarcane filter cakes present about 70% moisture, high organic matter content, calcium, potassium, magnesium, nitrogen and about 1.2 to 1.8% of phosphorus; due to these characteristics, the use of filter cake in the composition of organomineral fertilizers has been increasing since 1999, when the prices of mineral fertilizers reached high levels and environmental demands by consumers started to gain more space [7].

The evaluation of the quality and productivity of sugarcane indicated that the use of filter cake, enriched with soluble phosphate and applied in the furrow at planting, has the potential to partially replace mineral phosphate fertilization [8]. Organomineral fertilizers formulated with filter cake and sanitized sewage sludge (biosolid) positively affect parameters of growth and development of soybean plants and improve the activity of enzymes related to the protection of cell membranes [9]. The term biosolid is reserved for a stabilized byproduct; otherwise, the terms cake, sludge or solid are used [10,11]. The use of sewage sludge in agriculture began to be disciplined in Brazil by the CONAMA Resolution n° 375/2006 [12], establishing criteria for the safe use of this residue as fertilizer.

The benefits of biosolids as fertilizer have been observed in several studies with different crops. The incorporation of sewage sludge in the soil can improve its fertility and reduces the potential acidity proportionally to the dose applied [13]. This characteristic is probably due to the use of lime in the process of sanitization of the sewage sludge [14]. However, due to factors related to logistics and technologies of crop cultivation, the sources of organic matter as the filter cake and the sanitized sewage sludge began to present great commercial viability, especially when mixed with mineral fertilizers to compose organomineral fertilizers.

Organomineral fertilizers have distinctive characteristics; for example, phosphorus derived from these fertilizers was more rapidly available than from the mineral source (triple superphosphate) [15]. The organomineral fertilizers also provided a higher content of exchangeable potassium in the superficial soil layers in sugarcane cultivation, and are more efficient in all evaluated concentrations compared to exclusive mineral fertilization, which may replace the mineral fertilizers and provide up to 15% more efficiency in the production of sugarcane stalks [16].

In the case of sugarcane, the high productivity of stalks must be accompanied by physical-chemical parameters favorable to industry to guarantee high yields in the production of sugar and alcohol. However, the research still has much to contribute, justifying economically, environmentally and technically the use of organomineral fertilizers. Therefore, the objective of this study was to evaluate the chemical and technological characteristics of sugarcane fertilized with pelletized organomineral fertilizers based on filter cake and sanitized sewage sludge in two consecutive crop cycles.

## Material and methods

### Experimental area

The study was conducted in an experimental area of the *Companhia Mineira de Açúcar e Álcool* (CMAA), Vale do Tijuco Unity, located in Prata municipality, Minas Gerais state, Brazil, at 19° 29’ 59’’ S and 48° 28’ 26’’ W, and 780 meters a.s.l. The climate of the experimental area is tropical, semi-humid, classified according to Köppen as Aw - tropical dry winter [17]. Rains are concentrated between November and March, with an average annual rainfall of 1400 mm; the drought period concentrated between July and August. The average maximum temperature occurs in November to February ranging from 31 to 36 °C [18].

The experiment was installed upstream of a hill in a soil classified as Dystrophic Yellow Latosol (Oxisol) [19]. The soil is characterized as sandy soil, with 72% of sand, 18.5% of clay and 9.5% of silt. Soil sampling was performed at 0-0.2 and at 0.2-0.4 m depths and their chemical analyses are described in Table 1.

**Table 1.**
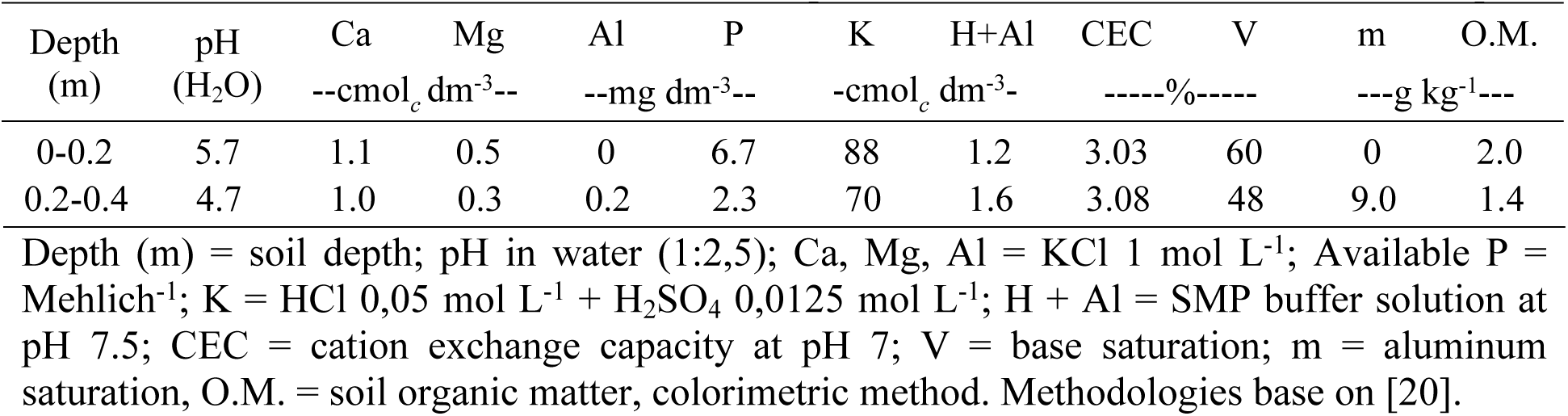
Soil chemical characterization of the experimental area at 0-0.2 and 0.2-0.4 m depth.

The soil of the experimental area received 2.4 t ha^−1^ of dolomitic lime, 1.5 t ha^−1^ of gypsum and 100 kg ha^−1^ of P_2_O_5_. Plowing with a moldboard plow (cutting width of 0.29 m spaced by 0.81 m) and leveling harrow (disks of 0.91 × 0.56 m) were used in the area. A degraded pasture occupied the experimental area for the past ten years.

### Organomineral fertilizers

At planting, 570 kg ha^−1^ formulation 04-21-07 was applied, plus 570 kg ha^−1^ of 07-00-28 + 0.7% of B at 150 days after planting [21]. Sanitized sewage sludge (biosolids) and sugarcane filter cake were used as sources of organic matter for the formulation of the organomineral fertilizers.

Sanitized sewage sludge was collected at the Municipal Department of Water and Sewage Treatment in Uberlandia, Brazil. The treatment in the station started by a railing and desander system (primary treatment), followed by a set of anaerobic reactors of the type UASB (upflow anaerobic sludge blanket reactor), systems with physicochemical polishing, centrifuges and geotextile systems destined to dewater the sewage sludge. The resulting material was centrifuged, separating the solids resulting in a product with 70% humidity and 30% solids. The wet sewage sludge went through chemical treatment incorporating 30% of hydrated lime [Ca(OH)_2_] and packaged in rectangular boxes of galvanized zinc (0.3×0.3×1 m). The structure was covered by a transparent canvas and exposed to sunlight for 15 days. Subsequently, the canvas was removed leaving to dry during 30 days and stabilizing at about 20% of moisture.

Following the laboratory report and the fertilization need for sugarcane planting, a pelletized organomineral was prepared: 39.3% of biosolid, 12.2% of crumbled KCl (58% K_2_O), 47% of crumbled monoammonium phosphate (12% N, 44% P_2_O_5_), and 1.5% of water. For the side dressing fertilization, the organomineral was prepared with 31% of biosolid, 15% of polymerized urea (45% N), 48.3% of crumbled KCl (58% K_2_O), 4.2% boric acid and 1.5% of water.

The pelletized organomineral fertilizer based on sugarcane filter cake used was produced by the Geociclo Biotechnology S/A (Uberlândia, Brazil). The composted filter cake received soluble mineral macronutrients (urea, monoammonium phosphate and KCl), boron as ulexite and micronutrients as oxisulfates. The resulting product received an organic polymer to produce the organomineral pellets. The organomineral pellets were about 3.9 mm diameter by 9.1 mm length.

The chemical characteristics of the sewage sludge and sugarcane filter cake usedare presented in Table 2.

**Table 2.** Chemical characterization of sewage sludge and sugarcane filter cake used in the composition of the organomineral.

| Attribute | Unity | Sewage sludge | Filter cake | Attribute | Unity | Sewage sludge | Filter cake |
| --- | --- | --- | --- | --- | --- | --- | --- |
| pH CaCl<sub>2</sub> | pH | 8.1 | 5.9 | Total mineral | % | 51 | 32 |
| Density | g cm<sup>-3</sup> | 0.66 | 0.65 | Boron | mg kg<sup>-1</sup> | 10 | < LQ |
| Total N | % | 0.99 | 1.79 | Sodium | mg kg<sup>-1</sup> | 201 | - |
| O.M. Total | % | 50 | 35 | Manganese | mg kg<sup>-1</sup> | 209 | 460 |
| Total carbon | % | 28 | 10 | Copper | mg kg<sup>-1</sup> | 135 | < LQ |
| C/N relation |  | 28/1 | 20/1 | Zinc | mg kg<sup>-1</sup> | 1042 | < LQ |
| Phosphorus | % | 2.80 | 2.25 | Iron | mg kg<sup>-1</sup> | 27,236 | 11,980 |
| Potassium | % | 0.30 | 0.30 | Cadmium | mg kg<sup>-1</sup> | 1.4 | 2.1 |
| Calcium | % | 8.25 | 2.43 | Mercury | mg kg<sup>-1</sup> | 0.7 | < LQ |
| Magnesium | % | 2.48 | 0.26 | Chrome | mg kg<sup>-1</sup> | 931 | 94.2 |
| Sulphur | % | 1.31 | 0.39 | Nickel | mg kg<sup>-1</sup> | 250 | - |

N - [N Total] = sulfuric digestion. P, K, Ca, Mg, S, Cu, Fe, Mn, Zn = nitro perchloric digestion. B = colorimetric azomethine-H. O.M. = organic matter. < LQ: lower than the limit of quantification. Methodologies based on [22].

The Geociclo Biotechnology S/A (Uberlândia, Brazil) also performed microbiological (total and thermotolerant coliforms) and chemical analysis in the organominerals. The total values of heavy metals: cadmium, chromium, nickel and lead, and the levels of total and thermotolerant coliforms were within the acceptable values determined by CONAMA Resolution n°. 375/2006.

### Experimental design

The experimental design was set as completely randomized in a 4×2+2 factorial scheme, being four doses of pelletized organomineral fertilizer, two organic sources (sewage sludge and filter cake), and two controls: positive control (exclusive mineral fertilization), and negative control (no fertilization), with four replications. The experimental plots measured 10 m long by 6 m wide, composed of five sugarcane rows spaced by 1.5 m from each other, totaling 60 m^2^. Each plot was separated by 3 m carriers. The useful area for the analysis was composed of three central lines of each plot dismissing the edge lines.

The sugarcane variety studied was ‘RB 92579’, a genotype of growth habit slightly decumbent, great biometric uniformity of stalk diameter and height, and medium tillering. This genotype came from the selection of the Northeast region of Brazil and has the characteristics of forming a long-lived sugarcane ratoon, high average productivity (average of five cuts), good closing of the lines and efficient planting and harvesting [23]. The planting was mechanized and carried out in the last week of May. In the planting furrows, 15 to 18 viable buds were sown per linear meter at a depth of 0.3 to 0.4 m. Simultaneously, the organomineral fertilizers were applied.

Eight treatments were combined being three levels of organominerals of two organic matter sources, plus two controls consisting of 100% of the recommended fertilization via mineral fertilizer and no fertilization; four replications were placed for each treatment. The levels of organomineral were 50, 100 and 150% of the recommended dose applied via pelletized organomineral fertilizer based on sanitized sewage sludge (SS50, SS100, SS150) or sugarcane filter cake (FK50, FK100, FK150).

The sugarcane was evaluated during two consecutive cuts (cycles): sugarcane of first year (sugarcane plant) and sugarcane of second year (first year of ratoon sugarcane). The control of insect pests (ants and termites) was performed with the application of 2.5 g ha^−1^ of fipronil in the furrow at planting, and for weed control, the herbicides hexazinone, diuron and MSMA were used in the respective doses 5, 3.2 and 3 L ha^−1^.

### Chemical and technological analysis

For the technological analyzes, ten stalks from the central lines of each experimental plot were collected. The stalks were clipped at the apical bud (breakpoint) and taken to the CMAA Laboratory for technological analyzes. The collections happened at 366 days after planting, and at 376 days after the first cut.

The quantity of stalks per hectare (ton ha^−1^), the sugarcane productivity (ton ha^−1^), and the quantity of sugar per hectare (ton ha^−1^) were estimated for each experimental plot. The processing was carried out according to the methodology based on sucrose content. After the disintegration and homogenization of the sugarcane stalks, an aliquot of 0.5 kg was subjected to a hydraulic press to extract the juice, which was used for the determinations chemical-technological [24]. The variables evaluated were:

Sugarcane *pol* (%): defined as the quantity of sucrose, in percentage, present in the sugarcane juice; *brix* of the juice (%): expresses the percentage (weight/weight) of soluble solids contained in a solution, i.e., measures the content of sucrose in the sugarcane juice; *purity* (%): determined by the relationship ‘pol/brix×100’; the higher the sugarcane purity, the better the quality of the juice to recover the sugar; and, *fiber* (%): represents the water-insoluble biomass in the sugarcane.

### Statistical analysis

Extreme values (*outliers*) of all variables evaluated were identified using boxplot graphs of residuals [25] generated by the software SPSS® Statistics. The outliers identified were calculated as a lost parcel to replace the extreme values.

The software SPSS Statistics® was also used to evaluate the ANOVA presumptions: Shapiro-Wilk’s [26] normality of residue distribution and Levene’s [27] homogeneity of variances, both at p > 0.01 probability.

After attendance of the ANOVA presumptions, the data were subjected to ANOVA (*F* test) to detect interactions between the factors (organic sources × doses) and differences among the levels in each factor (p < 0.05) [28]. When differences were detected appropriated comparisons were performed.

The additional treatments (mineral fertilization and no-fertilization) were compared to the factorial by the bilateral Dunnett’s test [29] (p < 0.05) using the software Assistat®.

The means of the factorial were compared by the Tukey’s test [30] (organic sources) at p < 0.05 probability, and by regression analysis (organomineral doses) using the software SISVAR®. The regression model was determined based on the significances of the coefficients (p < 0.05) and on the coefficient of determination (R^2^ > 70%). Sigma Plot® v. 12 software was used to compose the graphs of the organomineral doses.

The sugarcane crop yield variables between de first (366 days after planting in 2015) and second (376 days after the first harvest in 2016) sugarcane harvest were compared via a joint-analyses by the Tukey’s test [30].

## Results

The application of organomineral fertilizer based on sewage sludge at a dose of 100% (SS100) of the recommendation of fertilizer (100% mineral fertilizer), presented stalk production per hectare (SH) higher in the first year (Table 3).

**Table 3.**
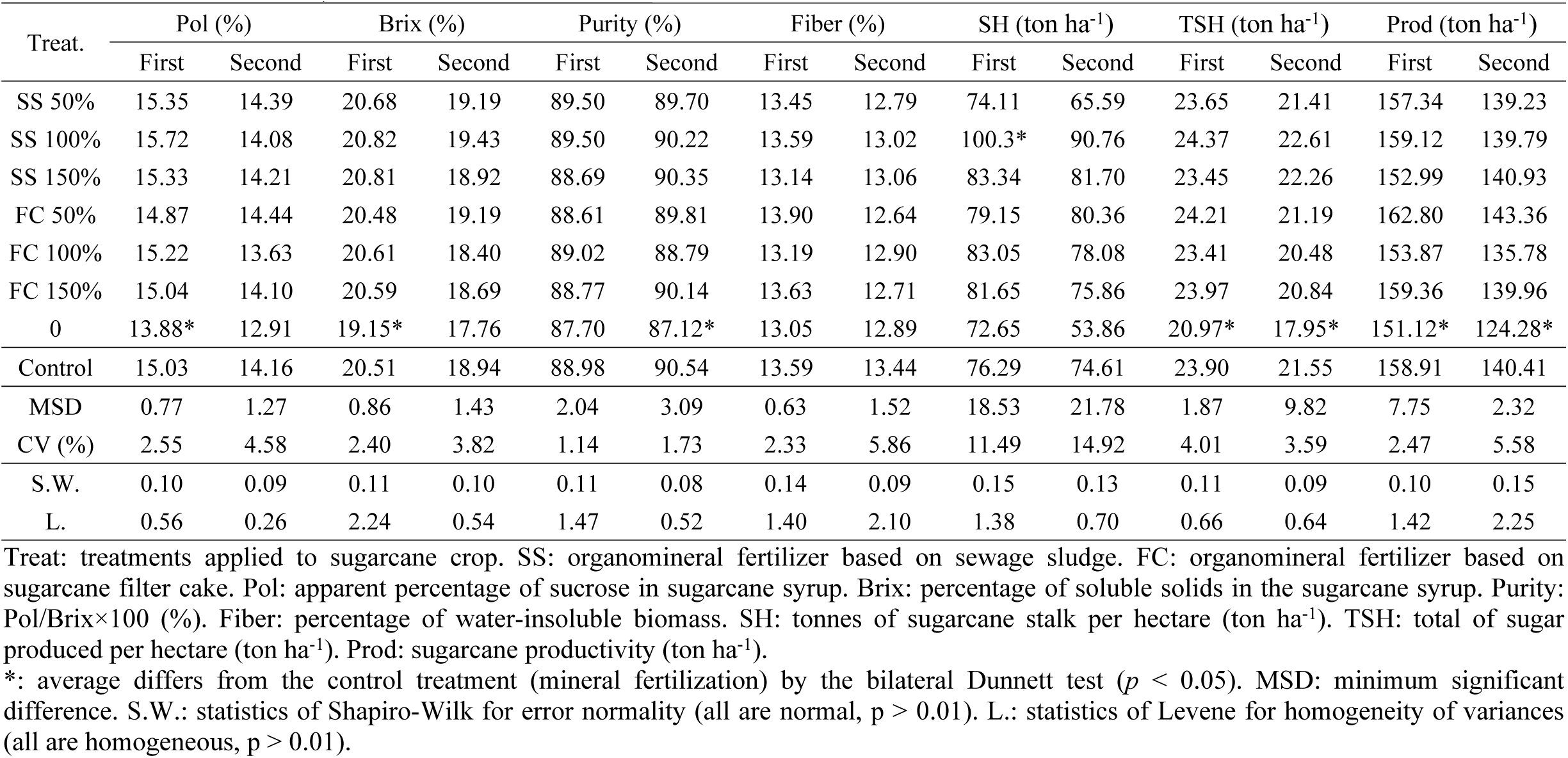
Average of chemical, technological and biometric variables of RB 92579 sugarcane cultivar (366 days after planting - first cut; 376 days after first harvest - second cut) for fertilization treatments including organominerals based on sewage sludge (SS) and sugarcane filter cake (FC).

| Treat. | Pol (%) |  | Brix (%) |  | Purity (%) |  | Fiber (%) |  | SH (ton ha<sup>-1</sup>) |  | TSH (ton ha<sup>-1</sup>) |  | Prod (ton ha<sup>-1</sup>) |  |
| --- | --- | --- | --- | --- | --- | --- | --- | --- | --- | --- | --- | --- | --- | --- |
|  | First | Second | First | Second | First | Second | First | Second | First | Second | First | Second | First | Second |
| SS 50% | 15.35 | 14.39 | 20.68 | 19.19 | 89.50 | 89.70 | 13.45 | 12.79 | 74.11 | 65.59 | 23.65 | 21.41 | 157.34 | 139.23 |
| SS 100% | 15.72 | 14.08 | 20.82 | 19.43 | 89.50 | 90.22 | 13.59 | 13.02 | 100.3* | 90.76 | 24.37 | 22.61 | 159.12 | 139.79 |
| SS 150% | 15.33 | 14.21 | 20.81 | 18.92 | 88.69 | 90.35 | 13.14 | 13.06 | 83.34 | 81.70 | 23.45 | 22.26 | 152.99 | 140.93 |
| FC 50% | 14.87 | 14.44 | 20.48 | 19.19 | 88.61 | 89.81 | 13.90 | 12.64 | 79.15 | 80.36 | 24.21 | 21.19 | 162.80 | 143.36 |
| FC 100% | 15.22 | 13.63 | 20.61 | 18.40 | 89.02 | 88.79 | 13.19 | 12.90 | 83.05 | 78.08 | 23.41 | 20.48 | 153.87 | 135.78 |
| FC 150% | 15.04 | 14.10 | 20.59 | 18.69 | 88.77 | 90.14 | 13.63 | 12.71 | 81.65 | 75.86 | 23.97 | 20.84 | 159.36 | 139.96 |
| 0 | 13.88* | 12.91 | 19.15* | 17.76 | 87.70 | 87.12* | 13.05 | 12.89 | 72.65 | 53.86 | 20.97* | 17.95* | 151.12* | 124.28* |
| Control | 15.03 | 14.16 | 20.51 | 18.94 | 88.98 | 90.54 | 13.59 | 13.44 | 76.29 | 74.61 | 23.90 | 21.55 | 158.91 | 140.41 |
| MSD | 0.77 | 1.27 | 0.86 | 1.43 | 2.04 | 3.09 | 0.63 | 1.52 | 18.53 | 21.78 | 1.87 | 9.82 | 7.75 | 2.32 |
| CV (%) | 2.55 | 4.58 | 2.40 | 3.82 | 1.14 | 1.73 | 2.33 | 5.86 | 11.49 | 14.92 | 4.01 | 3.59 | 2.47 | 5.58 |
| S.W. | 0.10 | 0.09 | 0.11 | 0.10 | 0.11 | 0.08 | 0.14 | 0.09 | 0.15 | 0.13 | 0.11 | 0.09 | 0.10 | 0.15 |
| L. | 0.56 | 0.26 | 2.24 | 0.54 | 1.47 | 0.52 | 1.40 | 2.10 | 1.38 | 0.70 | 0.66 | 0.64 | 1.42 | 2.25 |

Treat: treatments applied to sugarcane crop. SS: organomineral fertilizer based on sewage sludge. FC: organomineral fertilizer based on sugarcane filter cake. Pol: apparent percentage of sucrose in sugarcane syrup. Brix: percentage of soluble solids in the sugarcane syrup. Purity: Pol/Brix×100 (%). Fiber: percentage of water-insoluble biomass. SH: tonnes of sugarcane stalk per hectare (ton ha<sup>-1</sup>). TSH: total of sugar produced per hectare (ton ha<sup>-1</sup>). Prod: sugarcane productivity (ton ha<sup>-1</sup>).
\*: average differs from the control treatment (mineral fertilization) by the bilateral Dunnett test ( $p < 0.05$ ). MSD: minimum significant difference. S.W.: statistics of Shapiro-Wilk for error normality (all are normal, $p > 0.01$ ). L.: statistics of Levene for homogeneity of variances (all are homogeneous, $p > 0.01$ ).

The absence of fertilization (negative control) resulted in lower production of sugarcane per hectare and production of sugarcane than the positive control in both cycles. The application of SS100 reflected in a greater quantity of sugar produced per hectare (SH) in the second cycle, being 31.5% higher than that obtained in the negative control. Also, the negative control presented less apparent percentage of sucrose in sugarcane syrup (pol) and less percentage of soluble solids in the sugarcane syrup (Brix) in the first cycle of assessment when compared to treatment with mineral fertilizer (Table 3), while in the second cycle, only the sugarcane juice purity was reduced in the negative control.

In the first cycle, there was a significant interaction between the organic matter source used in the formulation of the organomineral fertilizers and the dose applied for sugarcane productivity. Only the sewage sludge source presented a significant (p<0.05) polynomial model for yield (Fig 1A), indicating that the highest yield (159.3 ton ha^−1^) observed was for a dose of 83.4% of referred organomineral fertilizer. The filter cake source (no polynomial adjustment detected) resulted in sugarcane productivity of 159.4 ton ha^−1^ for the 150% (FK150) dose of the recommendation of mineral fertilizer.

**Figure 1.**
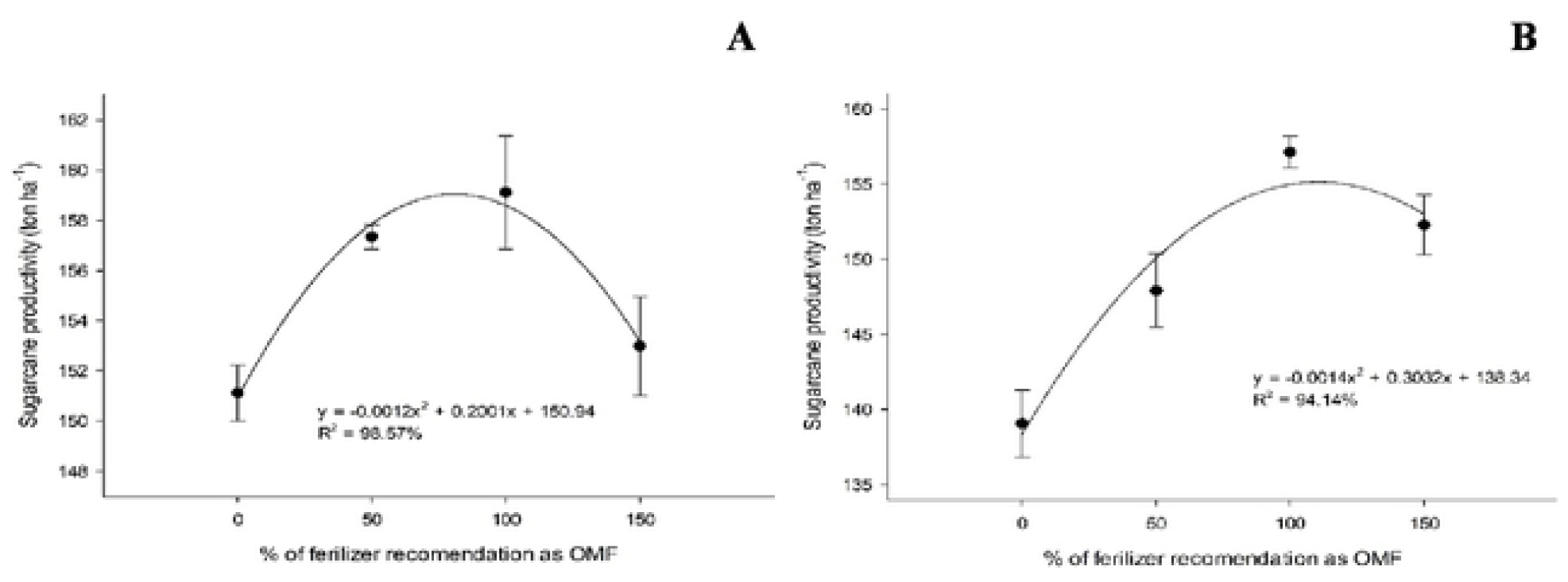
Sugarcane productivity (ton ha^−1^) under fertilizations with organomineral based on sewage sludge in the first (A) and second cut (B).

In the second cutting cycle, there was no interaction between sources and doses for sugarcane productivity; thus, the effect of the doses is independent of the organic matter source used (Fig 1B). The largest sugarcane production observed was 154.8 ton ha^−1^, obtained at a dose of 108.3% of organomineral fertilizer. The use of sewage sludge in the first cycle resulted in 17% savings in fertilization, with a contribution of 2.8% to the productivity compared to the second cut.

The highest yield in the first cut was observed when 50% of the recommended dose was used via organomineral fertilizer based on sugarcane filter cake (162.8 ton ha^−1^), and was approximately 7.73% higher than the negative control (151.1 ton ha^−1^). In this same dose, the organomineral fertilizer based on sugarcane filter cake resulted in high fiber production in the first cut (13.9%), compared to the fiber in the negative control (13.1%) (Table 3).

The greatest stalk productivity per hectare observed in the first cut was 86.7 ton ha^−1^ obtained at a dose of 110.2% of organomineral fertilizer regardless of the source of the organic matter used in the formulation of the organomineral. In the second cut, the greatest stalk productivity was 83.1 ton ha^−1^ at a dose of 108.7% of organomineral fertilizer (Fig 2). It is noticeable that the need for organomineral to express maximum stalk productivity was close between cuts, despite higher productivity (4.3% superior) in the first cut.

**Figure 2.**
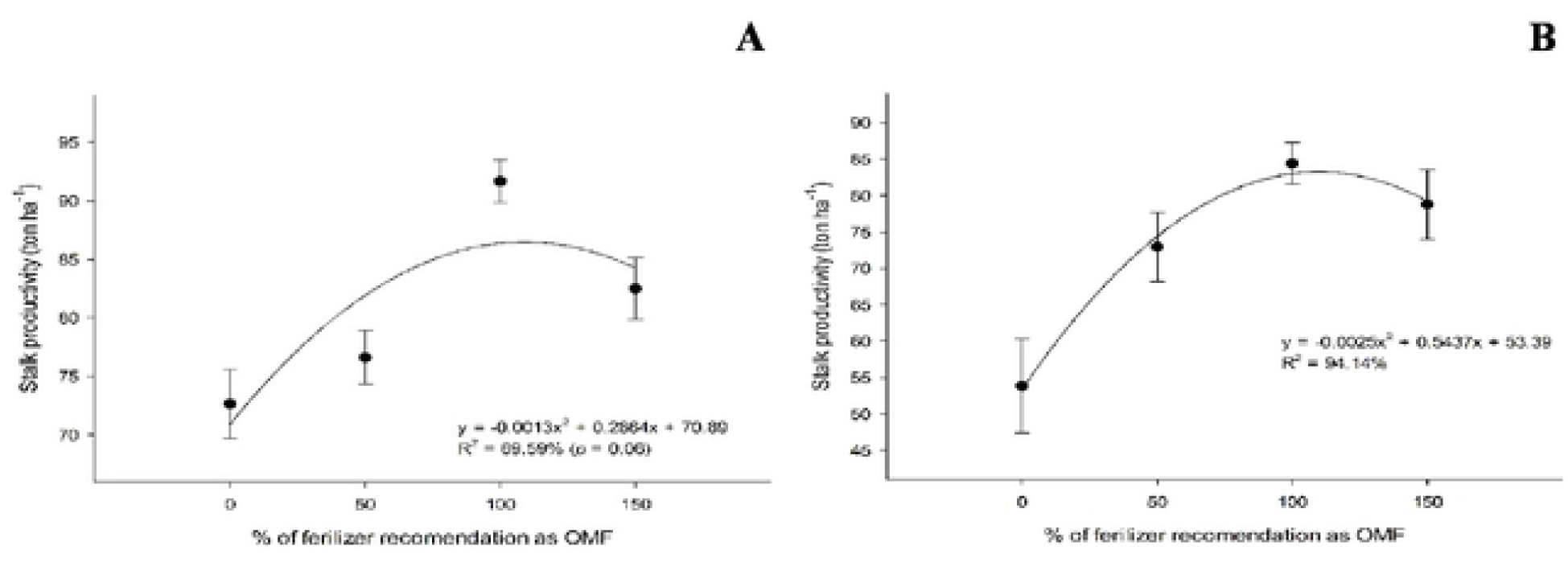
Stalk productivity (ton ha^−1^) under fertilizations with organomineral in the first (A) and second cut (B).

In the first cut, when the sewage sludge was the base for the organomineral fertilizer, the percentage of apparent sucrose (pol) was 2.16% higher than the pol observed when sugarcane filter cake was the base for the organomineral (Table 3). Similarly, in the second cut, when sewage sludge source was used, the quantity of sugarcane produced and the total amount of sugar per hectare was 4.19 and 4.68% higher than the sugarcane filter cake source, respectively (Table 4).

**Table 4.** Average of apparent sucrose (pol), sugarcane productivity and total of sugar produced for fertilization treatments including organominerals.

| Source | Pol (%) | TSH (ton ha<sup>-1</sup>) | Prod (ton ha<sup>-1</sup>) |
| --- | --- | --- | --- |
|  | -----First cut----- | -----Second cut----- |  |
| Filter cake | 14.75 b<sup>1</sup> | 20.11 b | 146.04 b |
| Sewage sludge | 15.07 a | 21.05 a | 152.15 a |
| MSD | 0.28 | 0.76 | 4.06 |
| CV (%) | 2.62 | 5.07 | 3.73 |
| S.W. | 0.10 | 0.09 | 0.15 |
| L. | 0.56 | 0.64 | 2.25 |

<sup>1</sup>Averages followed by distinct letters on the line differ from each other by the Tukey test a 0.05. S.W.: statistic of Shapiro-Wilk for error normality. L.: statistic of Levene for homogeneity of variances.

The largest percentage of apparent sucrose (pol) presented in the first and second cut, were 15.5 and 14.3%, respectively, for the doses of 104.1 and 102.1%, regardless of which of the sources was used in the fertilizer organomineral (Fig 3).

**Figure 3.**
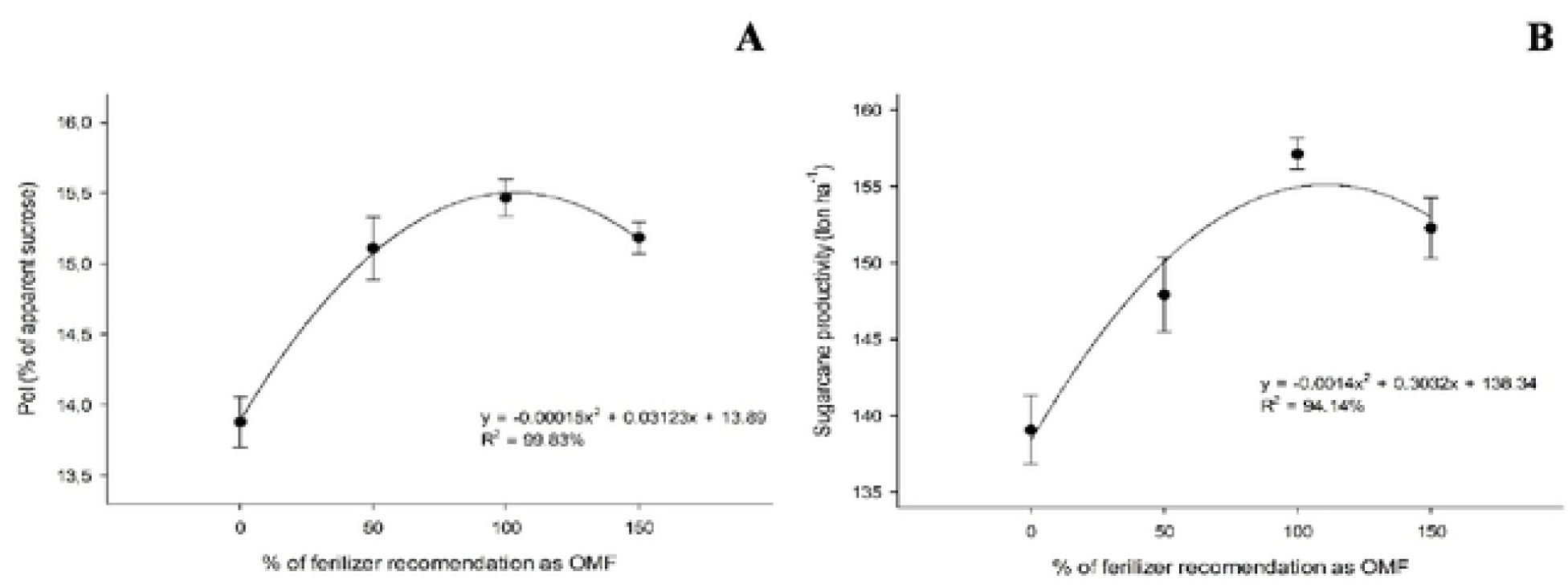
Sugarcane juice apparent sucrose (pol, %) under fertilizations with organomineral in the first (A) and second cut (B).

The percentage of soluble solids (brix) greatly varied with the organomineral fertilizer dose in both cycles, independently of the source of organic matter used. The largest percentage of soluble solids was 20.8% in the first cut and 19.2% in the second cut, respectively, for the doses of 104.3 and 96% of organominerals (Fig 4).

**Figure 4.**
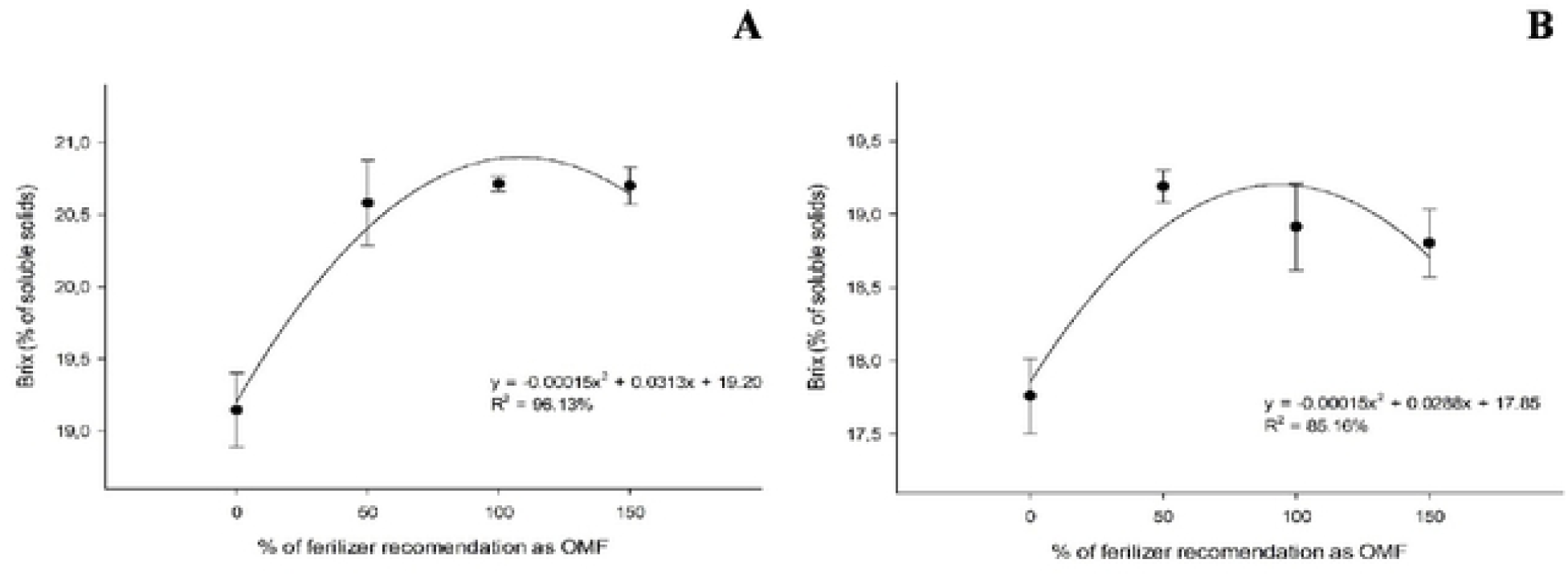
Percentage of soluble solids (brix, %) under fertilizations with organomineral in the first (A) and second cut (B).

In the first cut, the highest sugarcane juice purity (89.3%) was found for the organomineral fertilized dose of 91.6%, using sewage sludge or sugarcane filter cake. In the second cut, for each kilogram of organomineral fertilizer applied, using sewage sludge or sugarcane filter cake, there was an increase of 0.0182% in the purity of the sugarcane juice (Fig 5).

**Figure 5.**
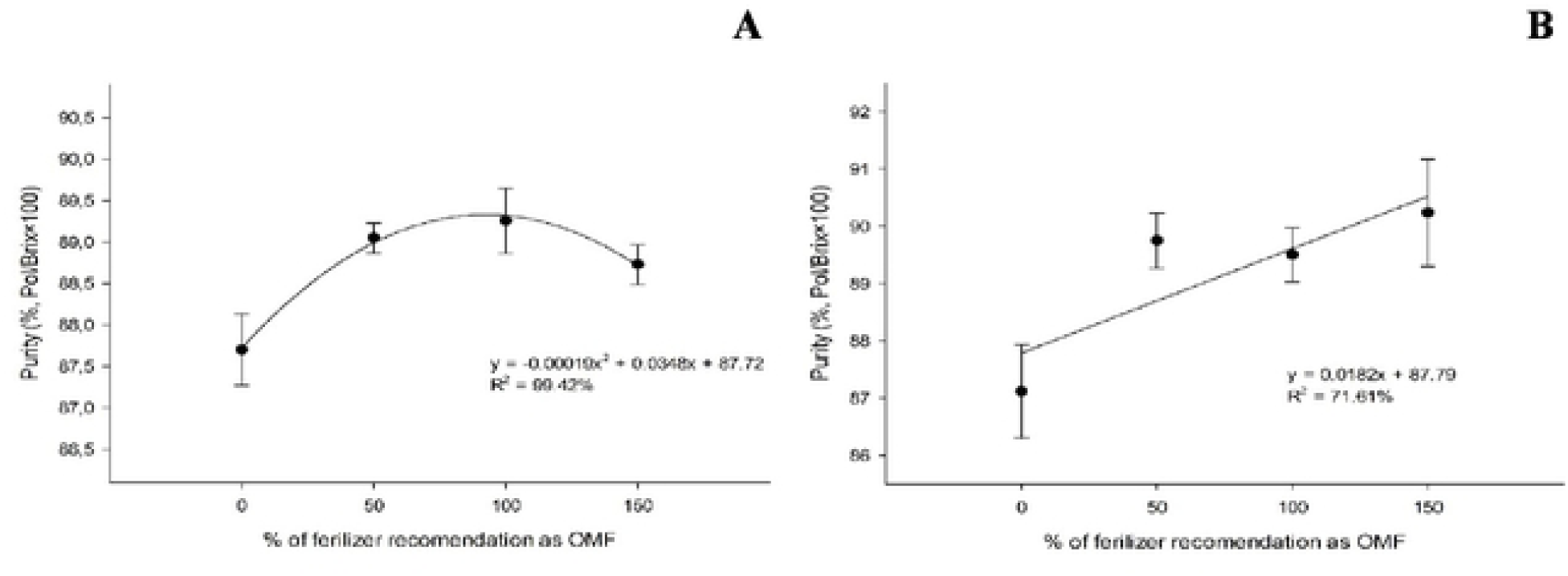
Sugarcane juice purity (%) under fertilizations with organomineral in the first (A) and second cut (B).

In the first cut, the highest fiber content observed was with the sewage sludge source (13.6%) obtained for the organomineral fertilized dose of 79.7% (Fig 6). There was no significant effect of doses and sources on fiber (%) in the second cut, being the average fiber found of 12.86%.

**Figure 6.**
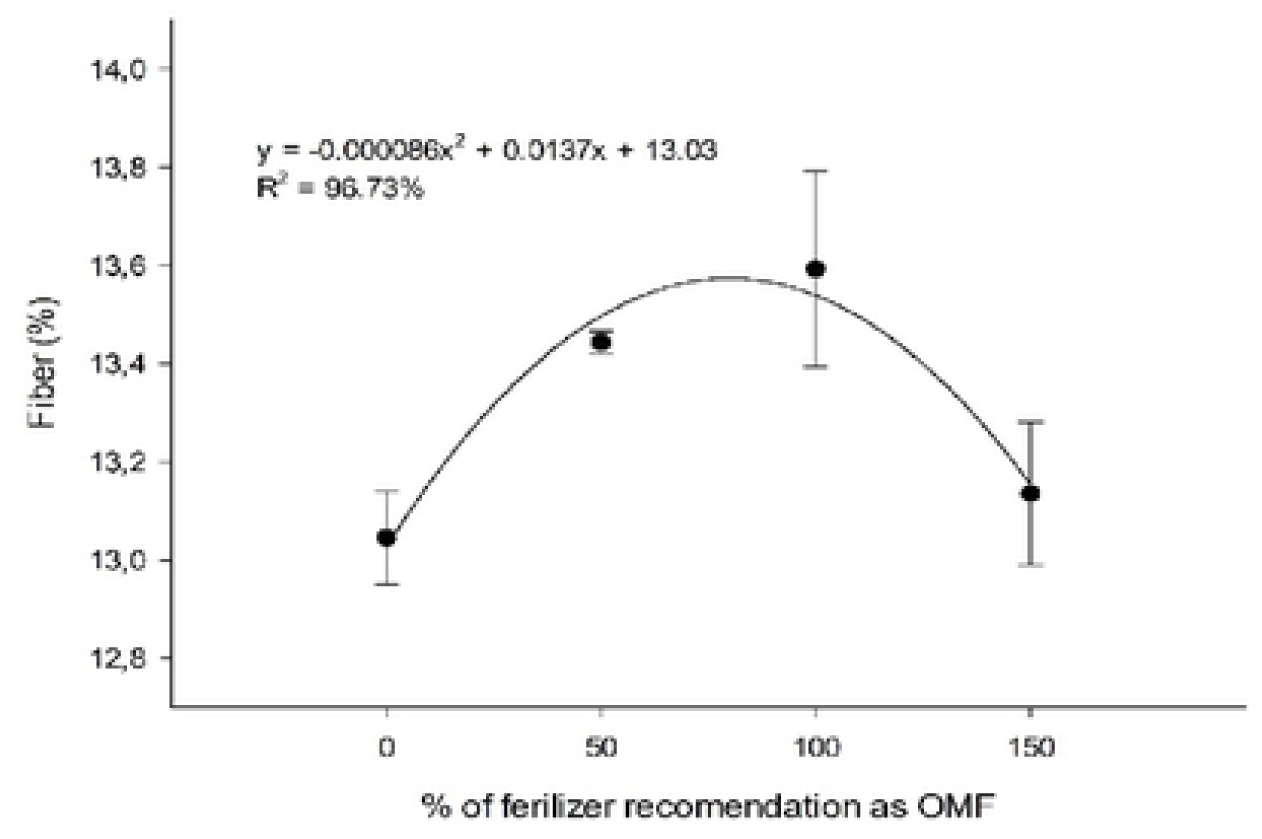
Sugarcane fiber (%) under fertilizations with organomineral based on sewage sludge in the first cut.

The largest quantity of sugar produced per hectare was 24.3 ton ha^−1^ in the first cut and 21.9 ton ha^−1^ in the second cut, respectively, for the doses of 102.3 and 106.5% of organominerals - using sewage sludge or sugarcane filter cake as organic sources (Fig 7).

**Figure 7.**
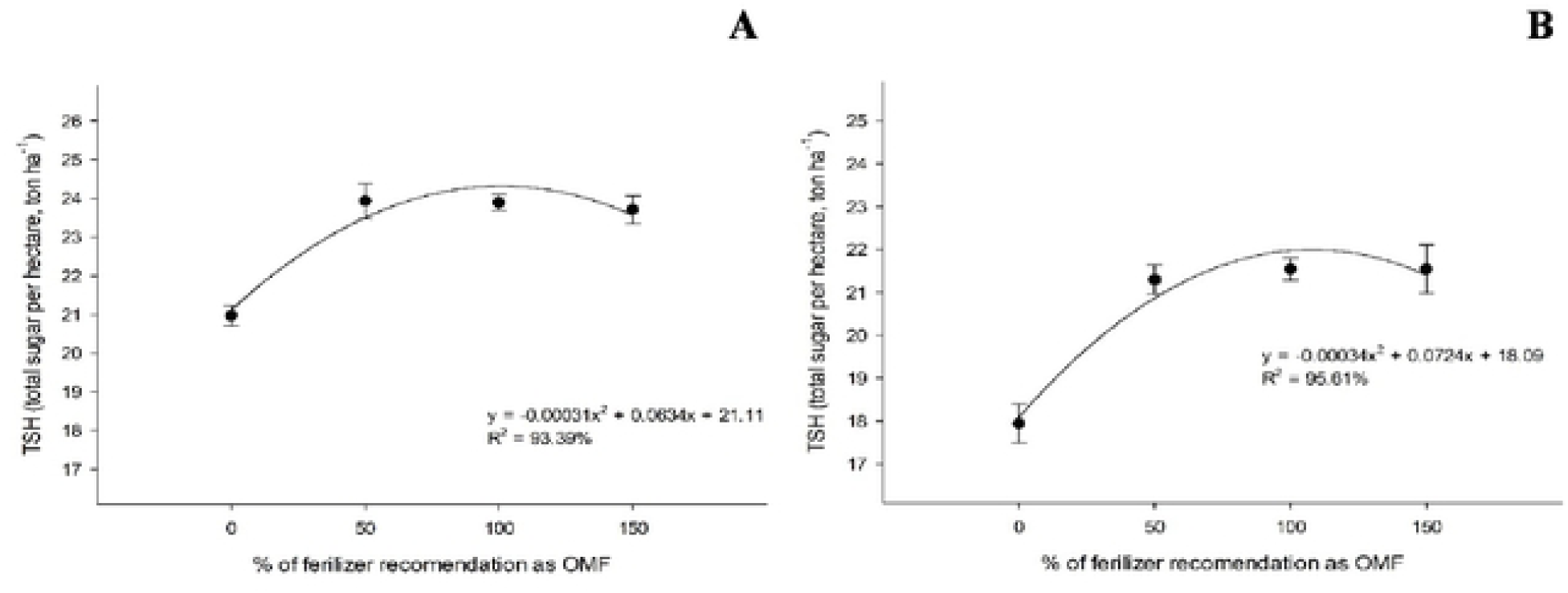
Total of sugar per hectare (TSH, ton ha^−1^) under fertilizations with organomineral based on sewage sludge in the first (A) and second cut (B).

The joint analysis between the first and second cuts indicated that the first harvest presented results greater than, or equal, to the second harvest, but never lower (Tables 5 and 6).

**Table 5.**
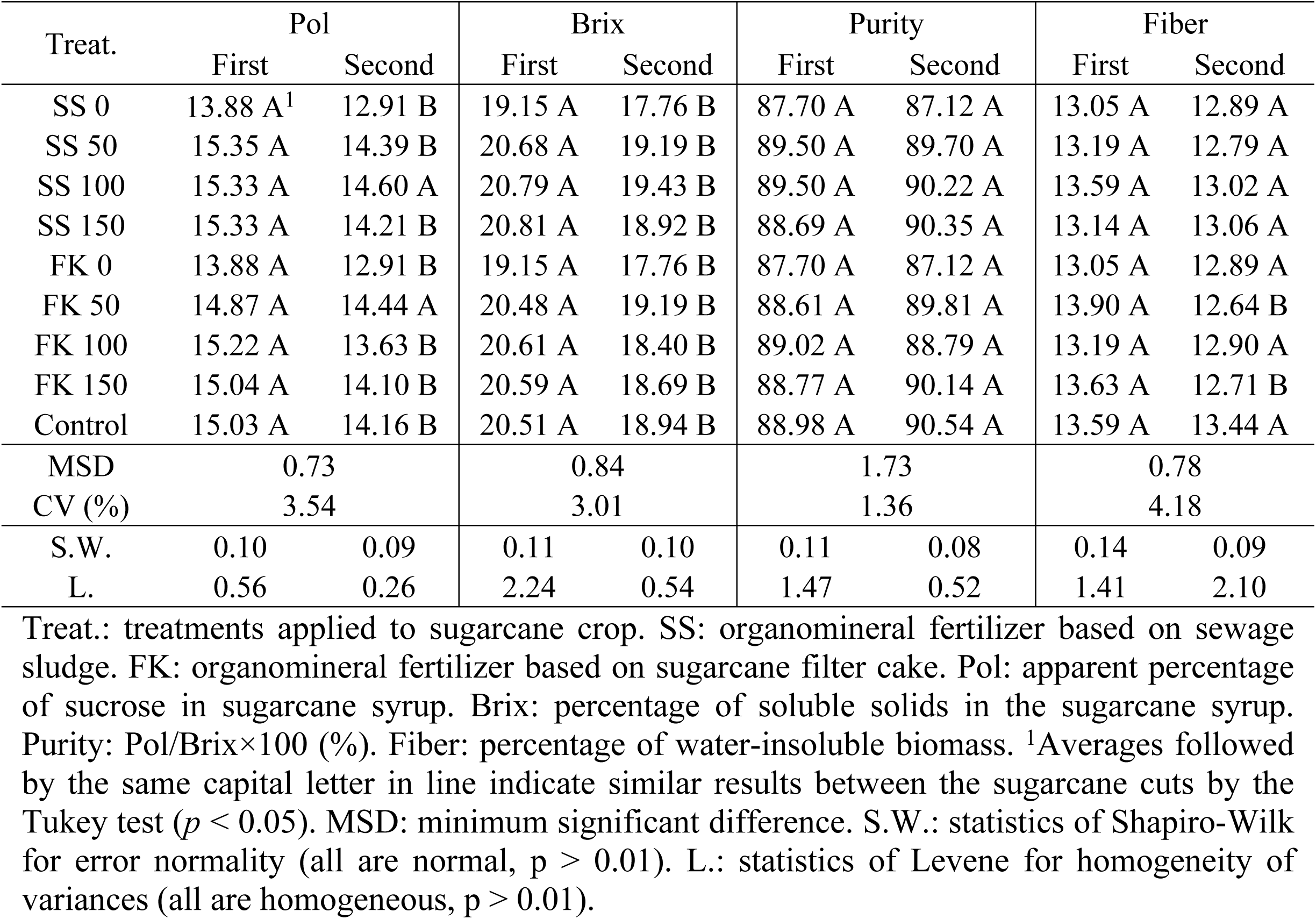
Joint-analyses of the sugarcane content of sucrose, soluble solid, purity and fiber variables between de first (366 days after planting) and second (376 days after the first harvest) sugarcane harvest.

| Treat. | Pol |  | Brix |  | Purity |  | Fiber |  |
| --- | --- | --- | --- | --- | --- | --- | --- | --- |
|  | First | Second | First | Second | First | Second | First | Second |
| SS 0 | 13.88 A<sup>1</sup> | 12.91 B | 19.15 A | 17.76 B | 87.70 A | 87.12 A | 13.05 A | 12.89 A |
| SS 50 | 15.35 A | 14.39 B | 20.68 A | 19.19 B | 89.50 A | 89.70 A | 13.19 A | 12.79 A |
| SS 100 | 15.33 A | 14.60 A | 20.79 A | 19.43 B | 89.50 A | 90.22 A | 13.59 A | 13.02 A |
| SS 150 | 15.33 A | 14.21 B | 20.81 A | 18.92 B | 88.69 A | 90.35 A | 13.14 A | 13.06 A |
| FK 0 | 13.88 A | 12.91 B | 19.15 A | 17.76 B | 87.70 A | 87.12 A | 13.05 A | 12.89 A |
| FK 50 | 14.87 A | 14.44 A | 20.48 A | 19.19 B | 88.61 A | 89.81 A | 13.90 A | 12.64 B |
| FK 100 | 15.22 A | 13.63 B | 20.61 A | 18.40 B | 89.02 A | 88.79 A | 13.19 A | 12.90 A |
| FK 150 | 15.04 A | 14.10 B | 20.59 A | 18.69 B | 88.77 A | 90.14 A | 13.63 A | 12.71 B |
| Control | 15.03 A | 14.16 B | 20.51 A | 18.94 B | 88.98 A | 90.54 A | 13.59 A | 13.44 A |
| MSD | 0.73 |  | 0.84 |  | 1.73 |  | 0.78 |  |
| CV (%) | 3.54 |  | 3.01 |  | 1.36 |  | 4.18 |  |
| S.W. | 0.10 | 0.09 | 0.11 | 0.10 | 0.11 | 0.08 | 0.14 | 0.09 |
| L. | 0.56 | 0.26 | 2.24 | 0.54 | 1.47 | 0.52 | 1.41 | 2.10 |

Treat.: treatments applied to sugarcane crop. SS: organomineral fertilizer based on sewage sludge. FK: organomineral fertilizer based on sugarcane filter cake. Pol: apparent percentage of sucrose in sugarcane syrup. Brix: percentage of soluble solids in the sugarcane syrup. Purity: Pol/Brix×100 (%). Fiber: percentage of water-insoluble biomass. <sup>1</sup>Averages followed by the same capital letter in line indicate similar results between the sugarcane cuts by the Tukey test ( $p < 0.05$ ). MSD: minimum significant difference. S.W.: statistics of Shapiro-Wilk for error normality (all are normal, $p > 0.01$ ). L.: statistics of Levene for homogeneity of variances (all are homogeneous, $p > 0.01$ ).

**Table 6.** Joint-analyses of the sugarcane crop yield variables between de first (366 days after planting) and second (376 days after the first harvest) sugarcane harvest.

| Treat. | SH |  | TSH |  | Prod |  |
| --- | --- | --- | --- | --- | --- | --- |
|  | First | Second | First | Second | First | Second |
| SS 0 | 72.66 A<sup>1</sup> | 53.86 B | 20.97 A | 17.95 B | 151.12 A | 139.06 A |
| SS 50 | 74.11 A | 65.59 A | 23.65 A | 21.41 B | 154.05 A | 148.86 A |
| SS 100 | 100.30 A | 90.75 A | 24.37 A | 24.00 A | 159.12 A | 163.98 A |
| SS 150 | 83.34 A | 81.70 A | 23.45 A | 22.26 A | 152.99 A | 156.71 A |
| FK 0 | 72.66 A | 53.86 B | 20.97 A | 17.95 B | 151.12 A | 139.06 A |
| FK 50 | 79.15 A | 80.36 A | 24.21 A | 21.19 B | 162.80 A | 146.98 B |
| FK 100 | 83.05 A | 78.08 A | 23.41 A | 20.48 B | 153.87 A | 150.26 A |
| FK 150 | 81.65 A | 75.86 A | 23.97 A | 20.84 B | 159.36 A | 147.85 B |
| Control | 76.29 A | 74.61 A | 23.90 A | 21.55 B | 158.91 A | 152.01 A |
| MSD | 14.40 |  | 1.49 |  | 7.32 |  |
| CV (%) | 13.22 |  | 4.77 |  | 3.36 |  |
| S.W. | 0.15 | 0.13 | 0.11 | 0.09 | 0.10 | 0.15 |
| L. | 1.38 | 0.69 | 0.66 | 0.64 | 1.42 | 2.25 |

Treat.: treatments applied to sugarcane crop. SS: organomineral fertilizer based on sewage sludge. FK: organomineral fertilizer based on sugarcane filter cake. SH: tonnes of sugarcane stalk per hectare (ton ha<sup>-1</sup>). TSH: total of sugar produced per hectare (ton ha<sup>-1</sup>). Prod: sugarcane productivity (ton ha<sup>-1</sup>). TRS: total of recovered sugar (kg ton<sup>-1</sup>). <sup>1</sup>Averages followed by the same capital letter in line indicate similar results between the sugarcane cuts by the Tukey test ( $p < 0.05$ ). MSD: minimum significant difference. S.W.: statistics of Shapiro-Wilk for error normality (all are normal, $p > 0.01$ ). L.: statistics of Levene for homogeneity of variances (all are homogeneous, $p > 0.01$ ).

## Discussion

The soil fertilization carried exclusively via mineral or via organomineral fertilizers did not diverge for the majority of the variables analyzed, indicating that organomineral fertilizers are a viable way of fertilizing sugarcane crop. The alternatives proposed in the present study, minimize the impacts arising from the successive application of mineral fertilizer sources, which are associated with the loss of soil biodiversity and the long-term dependence on external inputs [31, 32]. The positive environmental aspect of the use of sanitized sewage sludge and filter cake in organominerals in agriculture is safe and appropriate, according to the pre-treatment required for each source.

The organic fraction of organomineral fertilizer is an important aspect of these fertilizers. This fraction causes beneficial modifications that alter the dynamics of nutrient cycling in the soil-plant system [33]. Several studies also indicate organomineral fertilizers as viable substitutes for mineral fertilizers in many crops [8,34-38]. Regarding the soil dynamics of C and N, long-term field report demonstrated reductions in soil stocks of these nutrients in deep soil layers after the applications of large quantities of synthetic mineral fertilizers, which did not occur when the fertilizer source was an organomineral [39]. Organomineral fertilizers also reduce nitrogen (N-NH_3_) volatilization when compared to exclusively mineral sources [40].

Sugarcane filter cake also has considerable fractions of phosphorus, a critical element for agriculture in the Cerrado due to their low natural availability in the soils of this biome [41]. In addition to providing essential nutrients, organomineral fertilizers potentially increase the soil chemical quality due to the increase of the soil’s negative charges, extending the cation exchange capacity (CEC) and reducing the concentration of exchangeable aluminum (Al^3+^) [42-44]. The soil organic matter correlated with the regular applications of N and P, and with high soil CEC when sewage sludge was used as the source for organomineral fertilizer [45]. The monitoring of the environmental conditions and the orientation of farmers of this raw material (sewage sludge) on public policies is also essential to avoid heavy metals and biological contaminants during the reuse and management of this environmental liability [45].

Organomineral fertilizers based on sanitized sewage sludge can replace the mineral fertilization efficiently on maize [46] and sorghum [47] while improving soil fertility even in reduced doses and without heavy metal contamination. In soybean, reduced doses (25% less compared to mineral fertilizer) of pelletized organomineral fertilizer formulated with sanitized sewage sludge or sugarcane filter cake resulted in increments in plant height and stem diameter [9; 48]. In sewage sludge compost, the level of P present depends on the chemicals used in the process of the residue treatment to improve P concentration [49]. The release of nutrients to the environment occurs gradually in organomineral fertilizers, improving fertilizer efficiency [50]; thus, the P derived from organic residues constitutes a complementary strategy of supply of labile P to weathered soils [51].

The nitrogen:phosphorus (N:P) ratio of the organomineral with sewage sludge as the organic source is typically greater than that required by plants. In soils that receive sewage sludge organomineral plants accumulate more P than plants grown with only mineral fertilizers; the rate of P accumulation by the plants is influenced by the dynamics between climate and soil characteristics, especially in terms of the properties which determine the relationship between the proportion of adsorption and desorption of P from the soil [52]. The gradual P release is determined by the soil microbiota biodiversity and population present [53,54,4], and also by the residual effect of the fertilizer - organomineral fertilizers based on sugarcane filter cake can last for 4 years after its application [55,56].

The soil P concentrations in a study with maize [36] increased from 6 mg kg^−1^ (without fertilizer) to 24 mg kg^−1^ with the use of mineral fertilizer, and to 56 mg kg^−1^ with the use of organomineral fertilizer. The results found in that maize study were similar to what was found in the present study, in which the sugarcane production with organomineral fertilizer formulated with filter cake was comparable to the production where mineral fertilizer was applied. The root performance is also correlated to the effect of P in plant metabolism, being the presence of P fundamental for good rooting and sugarcane tillering, with a direct effect on final yield and sugar production [8]. Also, increments of P improve the apparent percentage of sucrose, the initial plant development and the P leaf content [57, 58]. In wheat, the application of filter cake also presented positive results, increasing grain yield with a tendency to interrupt the yield increase at doses greater than 60 ton ha^−1^ [59].

The organic fraction present in sanitized sewage sludge and in sugarcane filter cake organominerals contributes to improving soil aggregation, density, porosity, aeration, infiltration and the capacity of water retention [60]. Such factors are essential in the cultivation of sugarcane in months with low rainfall and in cultivations without irrigation [38,61]. In eucalyptus, sewage sludge compost increased the dry matter of the plants with a gain of 50% compared to the mineral fertilizer, indicating that this source of OM can replace the mineral fertilization [62]. The organomineral fertilizers have also the potential to significantly reduce the dependence on imports of mineral fertilizer sources of N, P and K. The reduction of costs of importation of P by a combination of features allows to increase economic productivity due to better use of energy and water [63], which increases the competitiveness of the agricultural sector. For example, the P obtained from sewage waste can recover the equivalent to 15 to 20% of the world demand phosphate rock [64].

The supplementation of the organic fraction with mineral elements can influence the use of both resources, with lower environmental impact and higher agronomic efficiency. Greater sugarcane yield was observed in the combination of 15 t ha^−1^ of filter cake and 350 kg ha^−1^ of NPK fertilizer [65, 66]. The effect of organic fertilizers in sugarcane using different organic sources (chicken bed, sugarcane filter cake, vinasse) associated with basic NPK fertilizer indicated that sugarcane filter cake provided the greatest amount of roots, which positively reflects in great productivity in the first and second cuts [60]. On the other hand, following the present study, no substantial increase in the production was found from the first to the second cut [38]. However, a study indicated a tendency of maintaining and even increase the productivity of stalks was regularly observed, since 13 clones, out of 25 clones, showed higher productivity with the application of organominerals [67].

The values observed for brix in the present study were higher than 18% for both harvests. The optimal value of brix is 18%, and it is an important variable for sugarcane yield since brix has a direct relationship with the sugar content and corresponds to 18 to 25% of the total of sugar [68]. The apparent percentage of sucrose (pol, %) is also important for the sugar industry which has an expectation of optimal value above 14% [69]. The values of sugarcane pol below 14% were observed in the treatment without fertilizer (negative control) in the first cut. The purity of the sugarcane juice represents the quality of the juice to recover its sugar, and the organomineral fertilizers (sewage sludge or filter cake) were similar to the mineral fertilizer, and all above 85% in both harvests. This value exceeds the ideal value estimated by the sugar industry [69], since the industry may refuse the receipt of shipments with purity below 75% [24]. Similarly, the fiber content (12 to 13%) in the first cut was within the ideal range considered adequate (10 to 13%) [69, 70], highlighting the efficiency of organominerals to provide nutrients to sugarcane crop adequately.

Favorable responses to the sugarcane cultivation have been observed at 120% of the fertilizer recommendation as organomineral, not differing from the 100% dose [71], a result that is similar to what was observed in this study. However, in consonance with the present study, little effect of the fertilization with organomineral in qualitative parameters is observed in the literature [72,73,5], highlighting the organominerals as efficient alternatives in the cultivation of sugarcane. The analysis of both harvests - the first and second harvests after the sugarcane planting - indicated what was expected for this crop culture, that the first cut is usually the most productive [74] since the responses of the first harvest were superior or similar to the second harvest.

This study allowed the observation that organomineral fertilizers can replace the exclusively mineral fertilization in sugarcane crop around the world, without negatively interfering in the quantity and quality of the attributes evaluated. The type of organic sources (sewage sludge or sugarcane filter cake) had little influence on the results observed; however, proper care must be taken on the collection, management and treatment of any organic source. In this way, a renewable resource, whose improper disposal can cause environmental impacts, generate favorable effects when added to the soil prepared for sugarcane

## Conclusions

The recommended organomineral dose to obtaining maximum quantitative and qualitative sugarcane results was between 102 and 109% of the regular recommendation, regardless of the organic source in the first sugarcane harvest.

In the second sugarcane harvest, sanitized sewage sludge source increase by 4.68 and 4.19% the total amount of sugar per hectare and the quantity of sugarcane produced compared to the sugarcane filter cake source.

Sanitized sewage sludge and sugarcane filter cake as sources for organominerals are viable alternatives and advantageous in economic and environmental terms for the cultivation of sugarcane.

## Acknowledgments

Coordination of Superior Level Staff Improvement (CAPES); National Council for Scientific and Technological Development (CNPq); Lutheran University of Brazil (ULBRA); Federal University of Uberlândia (UFU).

